# Fine-tuning protein language models on human spatial constraint improves variant effect prediction by reducing wild-type sequence bias

**DOI:** 10.1101/2025.10.15.682722

**Authors:** Gyasu Bajracharya, John A. Capra

## Abstract

Protein language models (PLMs) achieve state-of-the-art performance in predicting effects of missense variants, yet they do not explicitly consider variation within the human population. Here, we introduce <u>Hu</u>man <u>S</u>patial <u>C</u>onstraint (HuSC), a framework for quantifying intraspecies constraint on missense variants that integrates population-scale human genetic variation with 3D protein structures. We then fine-tune PLMs on HuSC scores. HuSC models the expected frequency of missense variation under neutral evolution and compares it to observed variation, accounting for both variation in mutational processes and 3D structural context. HuSC outperforms traditional inter– and intraspecies conservation metrics in predicting pathogenic variants. By focusing on intraspecies variation, HuSC reveals protein sites under human-specific constraint that cannot be captured by interspecies models. Integrating this intraspecies perspective into PLMs by fine-tuning on HuSC scores improves the prediction of variant fitness from deep mutational scans across diverse taxa and functional assay types. The improvement after fine-tuning comes largely from reducing bias toward wild-type sequences in regions that tolerate variation. Together, these results demonstrate that combining intraspecies constraint with cross-species PLMs improves their performance in variant-effect interpretation.

## 1 Introduction

The frequency of genetic differences within a population and between species is shaped by neutral processes and selection. Alleles that decrease organismal fitness tend to be lost over time, while fitness-increasing alleles are more likely to be retained. Comparing patterns of mutations and their frequencies to expectations under neutral evolution can highlight evolutionary constraints^1,2^. For example, sites in proteins essential for maintaining structure and function often exhibit little genetic variation due to strong evolutionary constraints. Quantifying these constraints both within and between species has provided valuable insights into proteins’ roles in disease and the selective pressures shaping their evolution.

Common methods for inferring evolutionary constraints between species rely on multiple sequence alignments (MSAs), where sequence conservation across species indicates constraint^3–6^. Recently, deep learning approaches such as protein language models (PLMs), trained on diverse protein sequences across the evolutionary tree, have expanded the ability to model constraints on sequence variation^7–10^. These models have achieved strong performance in predicting pathogenic variants identified in rare diseases and in estimating protein fitness from deep mutational scanning (DMS) data^11^. However, these methods do not inherently account for intraspecies sequence variation, which captures patterns of variation that reflect recent selective pressures over a much shorter period (thousands vs. millions of years). Additionally, the complexity of PLMs makes interpretation challenging; it is not always clear how they arrive at their predictions. This so-called ‘black-box’ nature may limit trust and broader adoption, especially in clinical settings.

The recent availability of large databases of human genetic variation across hundreds of thousands of individuals has enabled new strategies for quantifying evolutionary constraint within human populations^12–14^. Quantifying evolutionary constraint within the modern human population can provide relevant context for understanding the effects of variants on human disease. These methods have focused on quantifying constraints based on tolerance to missense or loss-of-function (LoF) variants at the protein or region level^15–20^. These methods have been further augmented by leveraging models of human protein 3D structures to consider the spatial context of variation^2,21,22^. These approaches become more powerful and comprehensive with the development of accurate computational methods for predicting 3D structure from sequence, but have not been competitive with interspecies conservation metrics in predicting site-specific variant effects.

Here, we unify these perspectives by improving the quantification of population-scale human genetic variation in the 3D structural context of proteins and integrating this signal into PLMs. First, we introduce the <u>Hu</u>man <u>S</u>patial <u>C</u>onstraint (HuSC) score, a framework that quantifies deviations in the frequency of missense variation within protein spatial contexts from expectations under neutral evolution. Then, we present an approach for incorporating these measurements of intraspecies constraint into PLMs through supervised fine-tuning. We show that HuSC outperforms traditional inter– and intraspecies conservation metrics in predicting pathogenicity. We also compare HuSC scores with interspecies conservation scores to identify protein sites that are more constrained within humans than across species. Furthermore, integrating HuSC into PLMs improves protein fitness predictions beyond the base model, highlighting the complementary roles of inter– and intra-species evolutionary constraints. By interpreting the sources of these performance gains, we find that fine-tuning with HuSC primarily recalibrates model confidence for residues in mutationally tolerant regions, leading to improved ranking of variants. Our results demonstrate that integrating intraspecies and interspecies constraints to capture both long-term evolutionary constraint and recent human-specific selection provides a more comprehensive view of a protein’s functional landscape and variant effects.

## 2 Results

### The Human Spatial Constraint (HuSC) framework maps evolutionary constraint in 3D protein structures

We developed the <u>Hu</u>man <u>S</u>patial <u>C</u>onstraint (HuSC) framework to quantify evolutionary constraint on 3D spatial regions of proteins in the human population. Our approach integrates human genetic variation data across hundreds of thousands of individuals with accurate models of protein 3D structures for ∼80% of human protein-coding genes. Many previous approaches, including our own COSMIS model, have quantified patterns of missense variation across human proteins. HuSC builds on these previous approaches by integrating strategies for considering protein 3D structural context, variable mutation rates at the nucleotide and protein scales, and the frequency distribution of observed variants.

The HuSC framework maps missense and synonymous variants and their minor allele frequencies (MAFs) across 141,456 individuals from gnomAD v2.1.1 into 16,259 3D protein structures from the AlphaFold Database (Fig. 1 & S1)^23,24^. To estimate the expected number and frequency of missense variants in a given 3D spatial region of a protein under neutral evolution, we built a permutation-based model that incorporates 3D spatial regions, local variation in the mutability of different nucleotide contexts, and global variation between proteins (Methods). Using this model, we calculate the expected frequency of missense variants in 3D spatial regions (e.g., spheres of varying radius) centered on each amino acid position in each protein.

**Figure 1:**
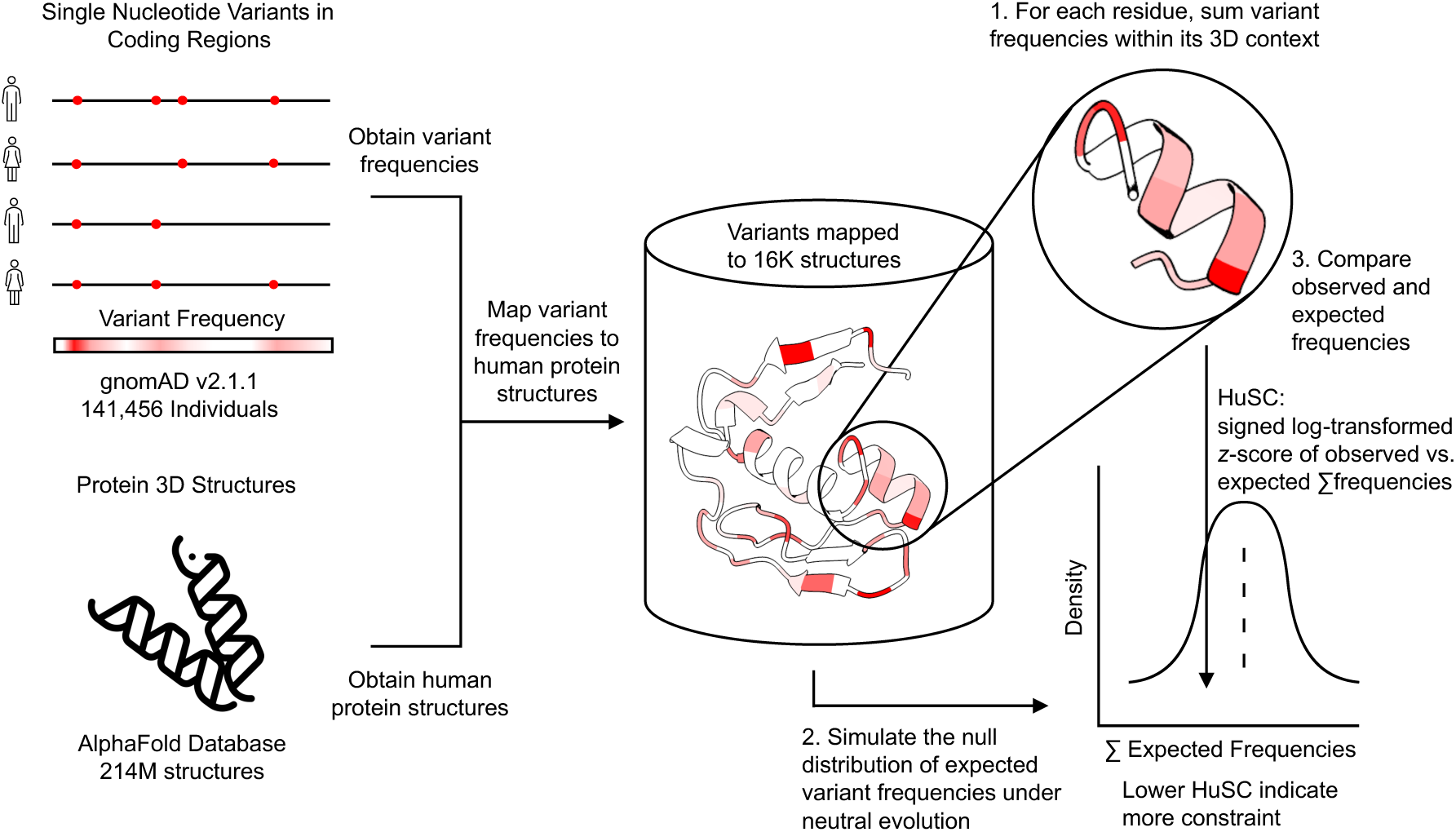
Schematic of the Human Spatial Constraint (HuSC) Framework. The HuSC framework integrates population genetics data with protein structure information by mapping single-nucleotide variant frequencies from hundreds of thousands of human genomes onto predicted 3D protein structures from the AlphaFold Database. For each protein residue, the framework aggregates the frequencies of variants observed within a 3D spatial region centered on the residue (e.g., an 8 Å sphere). The observed variant frequency within each 3D context for a residue is compared against a null distribution of total expected frequencies, generated using a mutation-spectrum-aware neutral model, to compute a signed log-transformed z-score (HuSC).

To compute the HuSC score, we compare the observed frequency of variants within a spatial region with the expected frequency under neutral evolution, as estimated by permutations (Methods). Specifically, for a given protein spatial region, we quantify its constraint by computing the observed frequency of variation in the spatial region minus the average of the expected frequency for the region over 10,000 permutations, divided by the standard deviation across permutations. We then log-transform this z-score to aid interpretability. Thus, lower HuSC scores signify a stronger constraint on a given 3D spatial region.

### Human Spatial Constraint (HuSC) scores highlight protein regions of functional importance

We applied the HuSC framework with an 8 Å radius context region to 8.8 million residues across 16,259 human proteins. Analysis of alternative context-region sizes revealed a trade-off between statistical power to detect constraint and spatial specificity, and 5–8 Å provided a good balance (Fig. S2).

As expected, the distribution of HuSC scores has a primary mode just below zero at - 0.37, reflecting that many protein-coding sites experience evolutionary constraint, where missense variation is less tolerated than expected under neutral evolution (Fig. 2A). It also has a broader positive mode extending into a long right tail at 0.41. This wider secondary mode corresponds to sites with more frequent variation than expected. The shorter tail for the negative scores (high constraint) is due to the fact that many of the most constrained regions do not have any variation, limiting the ability to differentiate among the most strongly constrained sites.

**Figure 2:**
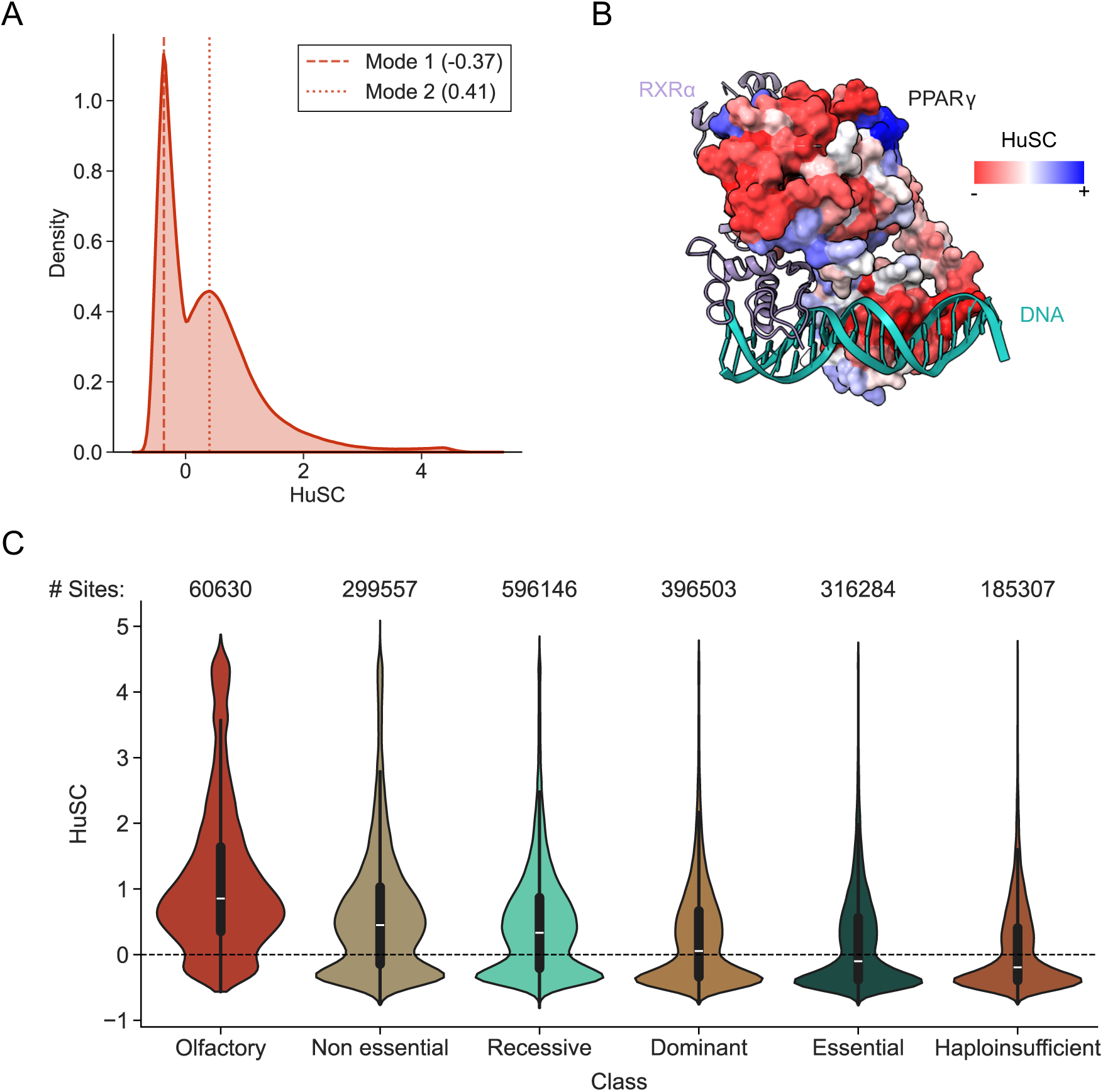
Human Spatial Constraint (HuSC) scores reveal functional constraint on protein 3D spatial regions. (A) Distribution of HuSC scores over 8.8 million residues across 16,259 human proteins. Negative scores indicate greater constraint, while positive scores indicate more variation than expected under neutral evolution. (B) Visualization of HuSC scores on the 3D crystal structure of Peroxisome Proliferator-Activated Receptor γ (PPARG) within the PPARγ–RXR–DNA complex (PDB: 3dzu). Residues are colored with negative HuSC values (red), indicating stronger constraints. The scores highlight strong constraints on the interfaces with RXRa (purple) and DNA (teal). (C) Violin plots of HuSC scores for all residues within proteins stratified by protein functional annotations. Proteins with essential functions (haploinsufficient and essential genes) exhibit higher constraint compared to proteins unlikely to influence fitness (olfactory receptors and non-essential genes). All pairwise comparisons between classes were statistically significant (Mann–Whitney U test, FDR-corrected p < 0.001).

As a qualitative example, we map HuSC scores on a crystal structure (PDB: 3DZU) of peroxisome proliferator-activated receptor γ (PPARG)^25^ (Fig. 2B). Mutations in PPARG are linked to numerous diseases, including diabetes^26^ and lipodystrophy^27,28^. PPARG functions as an obligate heterodimer with retinoid X receptor alpha (RXRα) and is capable of sequence-specific binding to DNA^29^. The HuSC scores reveal constraints that align with functionally important regions of PPARG. Residues in the DNA-binding and ligand-binding domains, especially at the binding interfaces, exhibit strong constraint. This visual example shows that HuSC scores are consistent with known functions and expected constraint on this protein.

Comparing HuSC scores across gene sets with known differences in evolutionary constraint and functional importance further supports that the scores behave as intended (Fig. 2C). Spatial regions in essential genes exhibit significantly lower HuSC scores than those in non-essential genes (median −0.1000 vs. 0.4510), consistent with stronger selective constraint. Similarly, dominant genes show lower HuSC scores than recessive genes (median 0.0560 vs. 0.3340). Haploinsufficient genes—where a single functional copy is insufficient for normal function—have the lowest HuSC scores (median −0.1930), reflecting their heightened sensitivity to variation. In contrast, olfactory receptor genes display the highest HuSC scores (median 0.8540), consistent with relaxed selective constraint in this gene family.

### HuSC outperforms other intraspecies and interspecies constraint metrics in pathogenicity prediction

HuSC quantifies intraspecies evolutionary constraint at the residue level based on patterns of human genetic variation. We therefore hypothesized that the constraint captured by HuSC would differ from that reflected by interspecies conservation metrics, which integrate evolutionary signal across species, and instead show greater similarity to other intraspecies residue– and gene-level constraint metrics.

To assess the similarity of HuSC scores compared to other methods, we computed Spearman correlations between HuSC and other constraint metrics across approximately 3.6 million residues in the human proteome (Fig. 3A). As expected, HuSC is most correlated with COSMIS (ρ = 0.61), a predecessor method that also integrates structural context with intraspecies genetic variation patterns. Its correlations with other intraspecies metrics, MTR3D and MTR, are lower (ρ = 0.34 and 0.35, respectively). HuSC shows similarly low correlations with intraspecies gene-level constraint metrics such as RVIS (ρ = 0.34), pLI (ρ = 0.36), and missense Z (ρ = 0.33). By contrast, its correlations with interspecies conservation metrics are even weaker—ConSurf (ρ = 0.29), phyloP (ρ = 0.26), GERP (ρ = 0.17), and PhastCons (ρ = 0.20)—consistent with these metrics capturing evolutionary conservation across species rather than population-level constraint. As expected, interspecies conservation metrics exhibit strong mutual correlations, particularly between phyloP and ConSurf (ρ = 0.79) and between PhastCons and GERP (ρ = 0.67).

**Figure 3.**
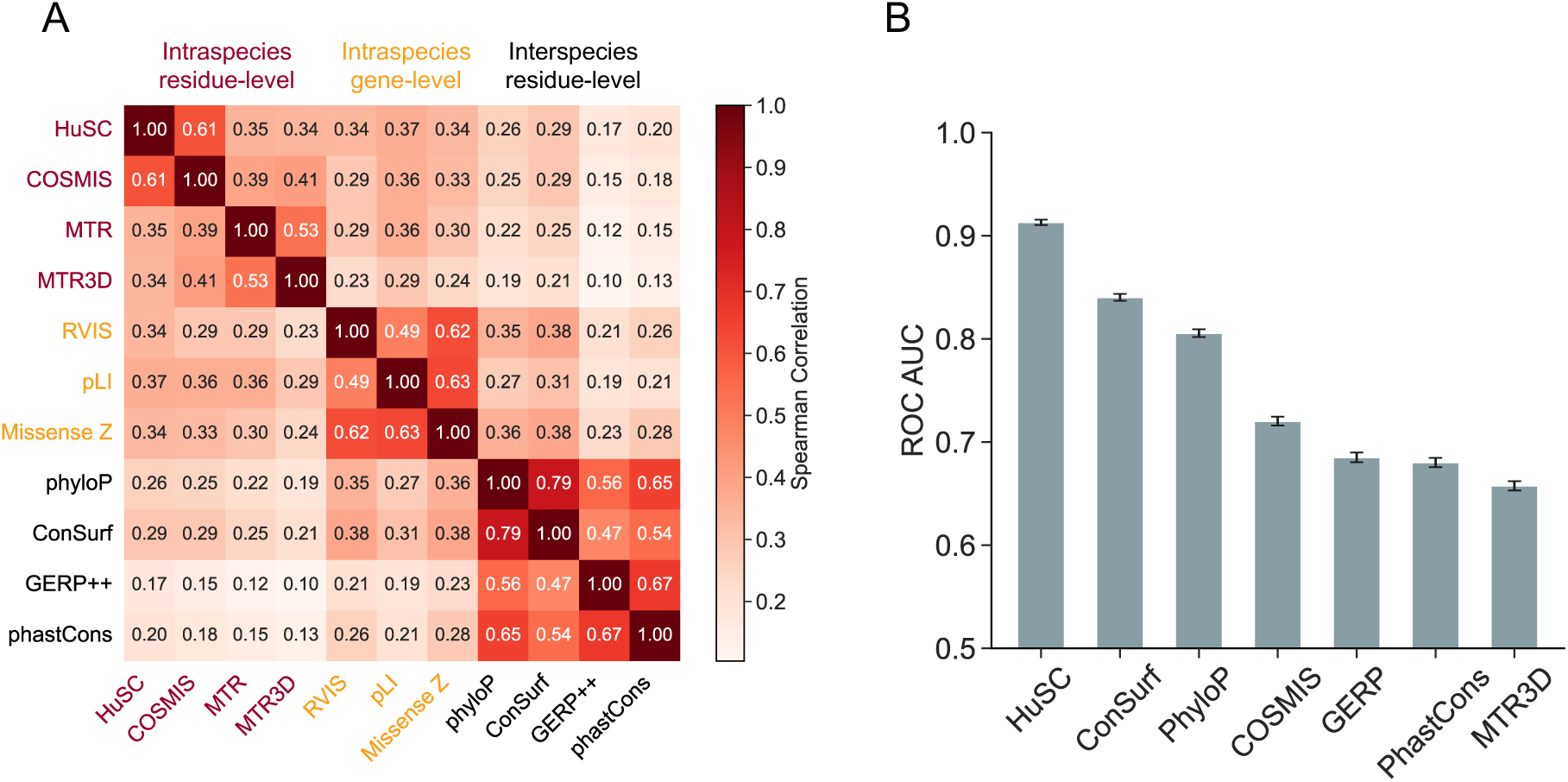
Comparison of HuSC with other intraspecies and interspecies conservation metrics. (A) Spearman correlation matrix comparing HuSC scores with other conservation metrics. HuSC is only modestly correlated with interspecies (phyloP, ConSurf, GERP++, phastCons), intraspecies residue-level (MTR3D, MTR), and protein-level constraint scores (RVIS pLI, Missense_z). The correlation was computed over 3.6 million sites across 12,107 proteins. (B) Receiver operating characteristic (ROC) area under the curve (AUC) values for classifying 6,416 pathogenic and 7,204 benign missense variants from ClinVar using HuSC and other conservation metrics. Error bars represent the standard deviation of AUCs calculated from 1,000 bootstrap resamples of the variant dataset. Pairwise comparisons of ROC AUCs between methods were performed using the Mann–Whitney U test with FDR correction for multiple testing, and HuSC achieved significantly higher performance than all other methods (FDR-corrected q < 0.05).

The HuSC framework does not directly quantify pathogenicity; rather, it quantifies constraint on missense variation in 3D spatial regions in human populations. Evolutionary constraint correlates with pathogenicity, since pathogenic variants are often found in highly constrained regions of the genome. As a result, constraint metrics are often used alone or in combination with other features to identify pathogenic variants. Thus, we compared HuSC’s performance at pathogenicity prediction against both intraspecies and interspecies constraint metrics.

We first evaluated the ability of different methods to distinguish pathogenic from benign missense variants in ClinVar using receiver operating characteristic (ROC) and precision–recall (PR) curve area under the curve (AUC) metrics (Fig. 3B & S3). The evaluation included 6,416 pathogenic and 7,204 benign variants curated from ClinVar.

HuSC outperforms all other conservation metrics in our comparison, achieving ROC and PR AUCs of 0.91 and 0.90, respectively. All pairwise comparisons of ROC and PR AUC between the evaluated scores were statistically significant (Mann–Whitney U test, FDR-corrected p < 0.05). The next best-performing methods are ConSurf (ROC = 0.84, PR = 0.79) and PhyloP (ROC = 0.81, PR = 0.75), both of which rely on interspecies conservation derived from multiple sequence alignments (MSAs). Among intraspecies constraint metrics, COSMIS (ROC = 0.72, PR = 0.74) and MTR3D (ROC = 0.66, PR = 0.68) perform comparably, whereas GERP (ROC = 0.69, PR = 0.60) and PhastCons (ROC = 0.68, PR = 0.60) show weaker performance. This does not indicate that HuSC is necessarily more accurate at quantifying constraint than these other metrics; rather, it suggests that HuSC’s quantification of constraint is more predictive of pathogenicity.

### Identification of human-specific constrained genes through comparison with interspecies conservation

Since interspecies and intraspecies conservation metrics capture distinct evolutionary signals (Fig. 3A), we leveraged their differences to identify regions that have evidence of constraint within humans but do not have strong conservation across species. The relationship between ConSurf scores (interspecies conservation) and HuSC scores shows moderate correspondence, with a Spearman correlation of *ρ* = 0.29, indicating that interspecies conservation explains only a limited fraction of the variance in human-specific constraint (Fig. 4A).

**Figure 4.**
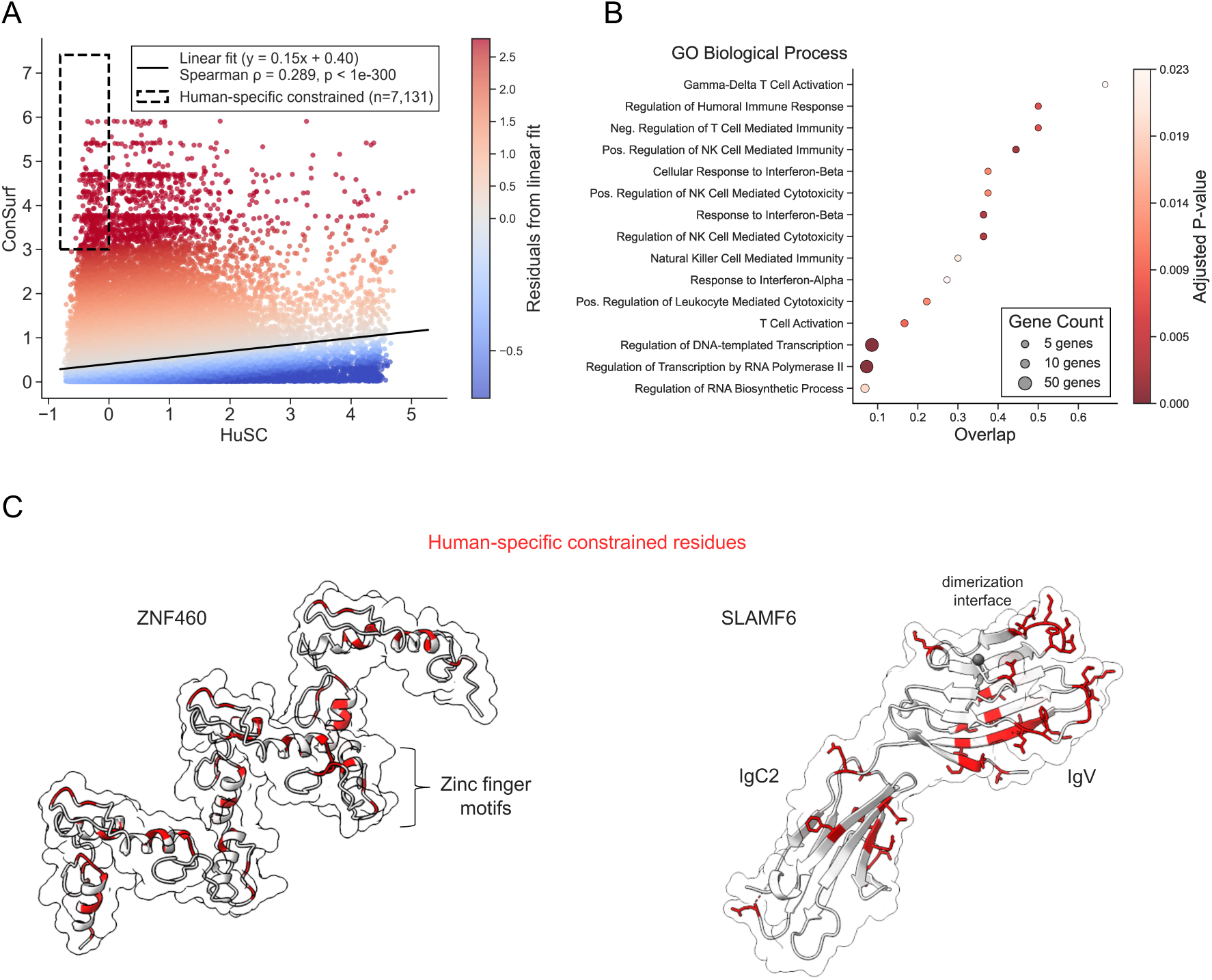
Identification and characterization of human-specific constrained genes. (A) Scatter plot showing the relationship between ConSurf scores (interspecies conservation) and HuSC scores across 2,125,803 residues from 5,643 genes. Spearman correlation is *ρ* = 0.289 (p < 1 × 10^−300^), with a linear fit of *y* = 0.15*x* + 0.40. The boxed region denotes the threshold used to define human-specific constrained (HuSC) residues (HuSC < 0 and ConSurf > 3). This region contains 7,137 residues corresponding to 1,093 unique genes. Point colors indicate residuals from the linear fit. (B) Dot plot showing Gene Ontology (GO) Biological Process enrichment for the 100 genes with the highest proportion of human-only constrained residues, using the GO Biological Process 2025 annotations. Dot color indicates the adjusted p-value, dot size reflects the number of genes annotated to each term, and ‘Overlap’ denotes the number of genes from each term present in the top 100 HuSC-enriched genes. (C) *(Left)* AlphaFold2-predicted structure of ZNF460 (AF-Q14592-F1), a representative gene from the human-specific constrained gene set, with human-only constrained residues highlighted in red. ZNF460 consists of multiple zinc finger motifs. *(Right)* Crystal structure of SLAMF6 (PDB: 2IF7), another representative HuSC-enriched gene, with human-only constrained residues highlighted in red. SLAMF6 comprises an IgV and an IgC2 domain, with the IgV domain forming the dimerization interface.

To identify residues under putative human-specific constraint, we applied a threshold of HuSC < 0 and ConSurf > 3. This criterion yielded 7,137 residues spanning 1,093 unique genes. We then ranked genes by the proportion of their residues falling within this thresholded region and selected the top 100 genes for functional enrichment analysis using the Gene Ontology (GO) Biological Process annotation dataset (Fig. 4B & S4). GO enrichment analysis revealed a strong overrepresentation of immune-related processes, including gamma-delta T cell activation, regulation of humoral immune response, negative regulation of T cell–mediated immunity, and positive regulation of natural killer cell–mediated immunity (adjusted p < 0.05). Among the proteins associated with positive regulation of natural killer cell–mediated immunity, Signaling Lymphocytic Activation Molecule Family Member 6 (SLAMF6) exhibited a high proportion of human-specific constrained residues. SLAMF6 is a homophilic receptor of the immunoglobulin superfamily, expressed on hematopoietic cells including T cells, natural killer (NK) cells, and B cells.

Structurally, SLAMF6 comprises an N-terminal IgV domain and a C-terminal IgC2 domain (Fig. 4C, right). Notably, the majority of human-specific constrained residues localize to the IgV-mediated dimerization interface, suggesting that human-specific constraint in SLAMF6 may reflect selective pressure on receptor–receptor interactions.

In addition to immune functions, we observed significant enrichment of transcription-related processes, most notably regulation of DNA-templated transcription and regulation of transcription by RNA polymerase II, which exhibited the most significant enrichment (p = 7.9 × 10^−22^ and 7.0 × 10^−16^, respectively). These transcription-associated terms were dominated by Krüppel-associated box (KRAB) domain–containing zinc finger proteins, a gene family known for rapid evolutionary turnover and pronounced species-specific diversification. One representative is ZNF460, which contains multiple zinc finger motifs (Fig. 4C, left). Human-specific constrained residues are distributed throughout these motifs, suggesting that selection within humans has shaped ZNF460-mediated DNA-binding and transcriptional regulation.

### Fine-tuning protein language models (PLMs) using Human Spatial Constraint (HuSC) improves protein fitness predictions

PLMs achieve state-of-the-art performance in predicting the effects of missense variants, yet they are not explicitly trained to model patterns of variation within the human population. We hypothesized that incorporating intraspecies spatial constraint information captured by HuSC would improve PLM performance on mutational effects prediction tasks. In this section, we demonstrate that HuSC scores are a useful complementary signal and illustrate an approach to incorporating them into PLMs via fine-tuning.

We fine-tuned ESM2^30^, a family of protein language models, using HuSC scores as supervision labels. We restricted training to the most constrained proteins and sites, where we expect intraspecies variation signals to diverge most strongly from the model’s pretrained interspecies evolutionary knowledge (Methods). Since we wanted ESM2 to retain its pre-trained knowledge of constraints across species, we applied low-rank adaptation (LoRA)^31^, which injects trainable rank-decomposition matrices, allowing the original model weights to be frozen and thereby mitigating catastrophic forgetting. For each protein sequence passed through ESM2, we first compute the log-likelihood ratio (LLR) for amino acid substitutions using the model’s log-probabilities. We applied a softmax over these LLRs to obtain the substitution probability distribution and calculated Shannon entropy of each position as a measure of local sequence constraint (Fig. 5A; Methods). The model was then trained to minimize a listwise ranking loss between predicted and observed HuSC values. We performed fine-tuning across four ESM2 model sizes with 8 M, 25 M, 150 M, and 650 M parameters (Fig. S5).

**Figure 5.**
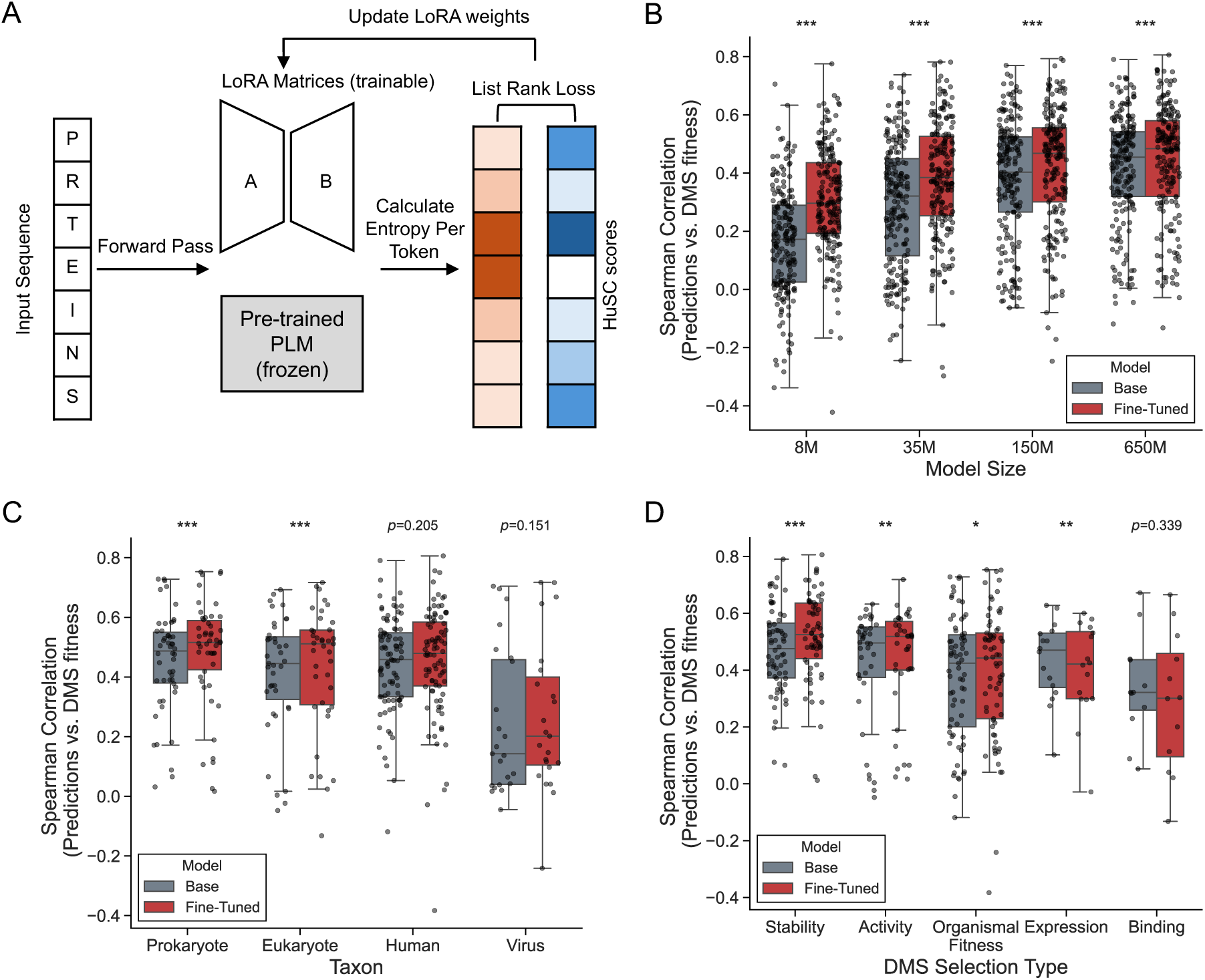
Fine-tuning ESM2 with HuSC scores improves protein fitness predictions. (A) Schematic of the fine-tuning strategy, where Human Spatial Constraint (HuSC) scores were used to fine-tune ESM2 models while preserving its pretrained evolutionary knowledge via low-rank adaptation (LoRA). (B) Comparison of Spearman correlations between model predictions and experimentally measured deep mutational scanning (DMS) scores for 201 proteins across ESM2 models of varying parameter sizes (8M, 35M, 150M, and 650M) before and after fine-tuning. Fine-tuned models show consistent significant performance gains across for all model sizes (Wilcoxon signed-rank test; *** p < 0.001). (C) Fine-tuning produces higher Spearman correlations with DMS data stratified by each protein’s taxon of origin: prokaryotic (n = 50, p = 1 × 10^−4^), eukaryotic (n = 39, p = 9 × 10^−4^), human (n = 89, p = 0.205), and viral (n = 23, p = 0.151). (D) Fine-tuned ESM2 650M achieves significant gains for stability (n = 66, p < 1 × 10^−4^), enzymatic activity (n = 38, p = 4.5 × 10^−3^), and organismal fitness (n = 69, p = 0.036), whereas expression (n = 16, p = 4.2 × 10^−3^) and binding (n = 12, p = 0.34) assays show mixed results.

To illustrate the benefits of our approach for another protein variant effect prediction task, we evaluated the fine-tuned models’ ability to predict variant effects on a range of protein properties, as estimated by deep mutational scanning (DMS) experiments. We used datasets from the ProteinGym benchmark, which comprises 250 standardized DMS assays spanning over 2.7 million mutated sequences across more than 200 protein families, covering a range of molecular functions, taxa, and depths of homologous sequence diversity^32^. Due to the fixed context window of the underlying model architecture, our analysis was restricted to the 201 DMS datasets corresponding to proteins of length ≤1022 amino acids. Fine-tuning ESM2 with HuSC scores improves missense variant effect prediction across all model sizes (Fig. 5B). For example, the median Spearman correlation between model predictions and DMS scores increased from 0.17 to 0.29 for the 8M model, 0.32 to 0.38 for the 35M model, 0.40 to 0.47 for the 150M model, and 0.45 to 0.48 for the 650M model (Wilcoxon signed-rank test, *p < 0.001* for all). As expected, the relative improvement from fine-tuning is smaller for larger models, but the gains remain statistically significant even for the largest (650M) model evaluated.

Notably, this improvement extends beyond human proteins. When stratifying DMS datasets by taxon, we observe significant increases in performance in eukaryotic (median ρ = 0.45 to 0.51, p = 9 × 10^−4^) and prokaryotic (0.49 to 0.52, p = 1 × 10^−4^) proteins after fine-tuning (Fig. 5C). This suggests that intraspecies spatial constraints capture fundamental aspects of protein function that generalize across evolutionary lineages. Furthermore, when categorizing DMS datasets by assay type, fine-tuning led to different amounts of improvement for different functional assays. The most significant improvements were for stability (0.48 to 0.53, p < 1 × 10^−4^), enzymatic activity (0.50 to 0.52, p = 4.5 × 10^−3^), and organismal fitness (0.42 to 0.44, p = 0.036), whereas expression and binding assays showed mixed or no improvement (Fig. 5D).

### Gains in fitness prediction arise from selective reduction of wild-type amino acid bias

Given the improvement in fitness prediction after fine-tuning ESM2, we sought to understand what the fine-tuned model was learning that led to these gains. We hypothesized that fine-tuning on human-specific residue-level constraints encourages the model to learn generalizable rules governing protein regions that tolerate variation.

To test this, we compared changes in overall performance across all substitutions, measured by Spearman correlation with changes in performance among the substitutions with the highest fitness values, measured by normalized discounted cumulative gain (NDCG) for the top 10% of variants in each DMS dataset (Fig. 6A). Improvements in NDCG for the top 10% of highest-fitness variants were strongly correlated with gains in Spearman correlation across datasets (Pearson *r* = 0.660, *p* = 1.53 × 10^−26^). Similar correlations were observed for NDCG at the bottom 10% of variants, indicating improved ranking of both highly fit and highly deleterious variants (Fig. S6A). Improvements in overall model performance, measured by Spearman correlation, were also very strongly correlated with gains in the ROC AUC for distinguishing fit vs. deleterious variants (Pearson *r* = 0.958, *p* = 1.71 × 10^−109^). This suggests robust performance gains for both fitness prediction and classification tasks (Fig 6B).

**Figure 6.**
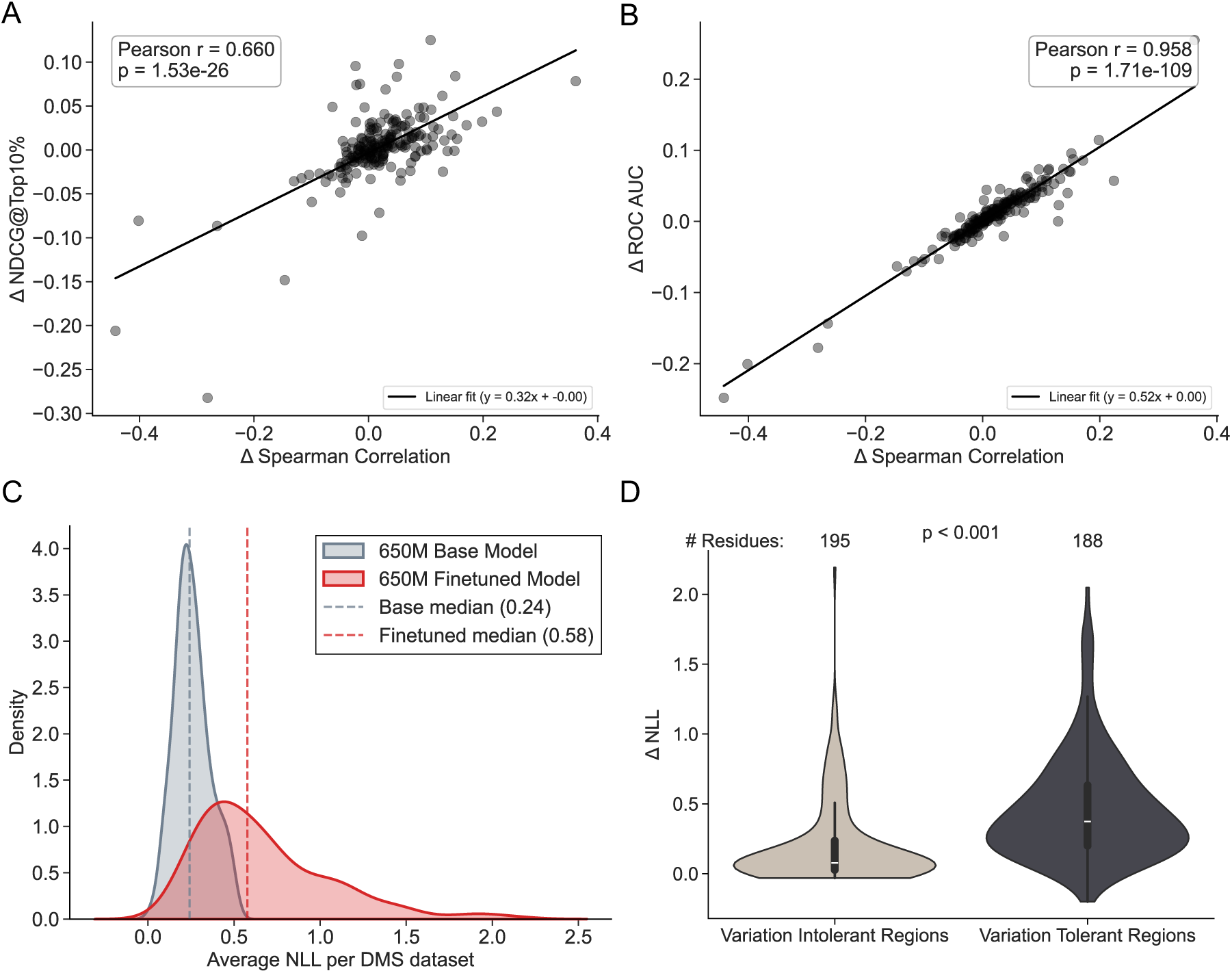
Changes in model performance metrics upon fine-tuning ESM2. (A) Scatter plots showing the relationship between changes in model performance across all substitutions in a protein and changes in performance on the 10% most fit substitutions across DMS datasets for 201 proteins from the ProteinGym benchmark. Performance on all protein substitutions is measured by Spearman correlation between model predictions and experimental fitness values. Performance on the most fit substitutions is quantified by normalized discounted cumulative gain (NDCG). Improvements in NDCG for the top 10% of highest-fitness substitutions are strongly correlated with gains in Spearman correlation across all substitutions (Pearson *r* = 0.660, *p* = 1.53 × 10^−26^; linear fit: y = 0.32x). (B) Scatter plot showing the relationship between changes in overall model performance (Δ Spearman correlation) and improvements in binary variant classification (Δ ROC AUC). Gains in ROC AUC are very strongly positively correlated with changes in ΔSpearman (Pearson *r* = 0.958, *p* = 1.71 × 10^−109^; linear fit: *y* = 0.52*x*). (C) Kernel density estimates of the average negative log-likelihood (NLL) of the wild-type protein sequence across DMS datasets for the base and fine-tuned models. Fine-tuning shifts the NLL distribution toward higher values, with the median increasing from 0.24 (base) to 0.58 (fine-tuned), indicating reduced confidence in wild-type sequence likelihoods. (D) Violin plots of changes in NLL (ΔNLL) upon fine-tuning for residues stratified by mutational tolerance. Residues belonging to variation-tolerant regions (top 10% of variants according to DMS fitness) exhibit significantly larger increases in NLL compared to variation-intolerant regions (bottom 10% of fit variants) (median ΔNLL: 0.373 vs. 0.077; Mann–Whitney U test p = 2.41 × 10^−20^).

To characterize how fine-tuning alters model behavior, we examined changes in the negative log-likelihood (NLL) of the wild-type protein sequence across DMS datasets (Fig. 6C). Fine-tuning consistently reduced the model’s confidence in wild-type amino acids (Fig. S6B). The average wild-type NLL increased substantially with the median across proteins rising from 0.24 in the base model to 0.58 after fine-tuning.

Notably, changes in NLL were not uniform across protein regions (Fig. 6D & S6C). When residues were stratified into variation-intolerant (bottom 10% of variants according to DMS fitness) and variation-tolerant regions (top 10% of variants by DMS fitness), variation-tolerant regions exhibited significantly larger increases in NLL after fine tuning (median ΔNLL: 0.373 vs. 0.077; Mann–Whitney U test, p = 2.74 × 10^−20^). This indicates that fine-tuning preferentially reduces overconfidence in the wild-type amino acid for regions that tolerate multiple amino acid substitutions.

Together, these results suggest that fine-tuning primarily improves fitness prediction by recalibrating the model’s confidence for wild-type residues in mutationally tolerant regions.

## 3 Discussion

Quantifying mutational constraint within the human population can provide valuable insights into the functional importance of genomic regions and the fitness of different amino acid substitutions. While deep learning models such as protein language models (PLMs) achieve state-of-the-art performance in predicting effects of missense variants^8,9^, they are primarily trained on representative sequences from across the tree of life and do not explicitly model within-species population variation. To address these limitations, we developed the Human Spatial Constraint (HuSC), a new metric that integrates large-scale human genome and exome sequencing data with protein structure predictions to quantify constraint in 3D spatial regions. HuSC outperforms existing population-variation-based metrics at predicting pathogenic variants and captures signals that are complementary to interspecies conservation metrics. Furthermore, fine-tuning PLMs with HuSC scores improves model performance and reveals that incorporating within-human constraints recalibrates confidence in wild-type residues, particularly in mutationally tolerant regions.

The HuSC framework quantifies evolutionary constraint by integrating a protein’s 3D spatial regions, allele frequency information, and background mutation rates. Since spatially proximal residues often share similar selective pressures, incorporating 3D context provides a more biologically meaningful representation of constraint across the proteome. It also increases power by considering variation patterns across multiple sites. Previous methods, including our earlier approach (COSMIS), primarily considered the presence or absence of variants and thus overlooked valuable information contained in variant frequencies^2^. Since allele frequency reflects the relative tolerance of a site to mutation, we hypothesized that incorporating both the occurrence and prevalence of variants contribute to more accurate constraint estimation. Consistent with expectations, HuSC scores reveal stronger constraint in essential and disease-associated genes. HuSC scores also outperform other conservation metrics in classifying pathogenic variants. Together, these findings support our hypothesis that combining structural context with allele frequency information yields a more informative and robust measure of mutational constraint.

HuSC predominantly reflects selective pressures over the past few hundred thousand years of human evolution rather than evolutionary conservation between species over hundreds of millions of years. Comparing within and between species constraint metrics revealed considerable differences, suggesting that intraspecies constraint captures a signal that is distinct from interspecies conservation metrics. Leveraging this distinction, we identified residues under putative human-specific constraint (low HuSC scores but relatively high interspecies conservation), and observed strong functional enrichment in immune-related processes. Among these, SLAMF6 exhibited a particularly high proportion of human-specific constrained residues. SLAMF6 is a homotypic receptor expressed on T cells, natural killer cells, B cells, and dendritic cells, and has recently been shown to function as an inhibitor of T cell activation, in both humans and mice, with this inhibitory activity mediated by homotypic *cis* interactions^33–35^. Notably, the majority of human-specific constrained residues in SLAMF6 localize to the IgV domain that forms the homodimerization interface. This suggests the human-specific constrained residues within SLAMF6 may modulate this receptor–receptor interaction in a lineage-specific manner.

We also observed strong enrichment of transcription-related processes mediated by KRAB zinc finger proteins. These proteins are characterized by an N-terminal Krüppel-associated box (KRAB) domain and a C-terminal array of two to more than forty zinc finger motifs with sequence-specific DNA-binding potential^36^. As a result, KRAB-associated zinc finger (KRAB-ZNF) proteins function as potent transcriptional repressors^37–39^. Proteins combining KRAB and zinc finger domains have expanded rapidly in mammals to include hundreds of members^40–42^. The rapid expansion and evolutionary turnover of KRAB-ZNF genes are thought to be driven largely by changes in the transposable element (TE) landscape of their hosts, giving rise to highly species-specific repertoires^43,44^. In this context, the high proportion of human-specific constrained residues observed across zinc finger motifs in proteins such as ZNF460 is consistent with ongoing lineage-specific selection acting on DNA-binding interfaces. Recent work studying the evolution of KRAB-ZNFs provides evidence for ongoing evolution of KRAB-ZNF proteins in modern humans^45^.

To demonstrate the promise of our observation that inter– and intraspecies constraint metrics capture complementary signals, we present an approach that integrates HuSC scores into protein language models (PLMs), which are typically not trained to capture patterns of sequence variation within human populations. To incorporate this information, we used HuSC scores as a supervision signal to fine-tune PLMs without altering the model architecture. We focused fine-tuning on the most constrained proteins and sites, where we expected the strongest complementary signal to emerge. We applied LoRA-based fine-tuning, making this approach easily transferable to other PLMs. Performance gains diminished with increasing model size, suggesting that larger models may encode aspects of population-level constraint derived from diverse evolutionary data; however, we propose that direct integration of within-species variation is a more efficient way to achieve these benefits. Although we anticipated improvements would be confined to human proteins, the benefits generalized across taxa, likely reflecting conserved patterns of functional constraint that are more easily captured in population-scale variation data. Improvements were most pronounced for functional assays related to stability, organismal fitness, and activity, which may be more directly linked to evolutionary constraint than binding or expression.

To understand the source of the observed performance gains, we examined how fine-tuning with HuSC alters the PLM. Interestingly, fine-tuning improved the prioritization of both the most fit and the most deleterious variants, even though training was limited to constrained HuSC scores. Moreover, discrimination between the most fit and deleterious variants also improved, suggesting that overall performance gains arise from changes in how models represent mutational tolerance as well as increased sensitivity to mutational constraint. Consistent with this interpretation, fine-tuning increased the model’s negative log-likelihood for wild-type residues, indicating reduced weight on the reference sequence. This recalibration was most pronounced in regions that tolerate substantial variation, where overconfidence in wild-type residues is most likely to obscure the relative fitness of alternative amino acids. Together, these findings suggest that incorporating within-human constraint encourages PLMs to learn generalizable principles of mutational tolerance, improving fitness prediction by correcting overconfident priors in permissive regions rather than by sharpening predictions at highly constrained sites.

Our work has several limitations. First, HuSC currently applies a linear weighting of allele frequencies, even though the relationship between allele frequency and variant effect size is inherently nonlinear. Incorporating a nonlinear weighting scheme (such as logarithmic scaling) could enhance sensitivity to low-frequency variants while reducing the influence of high-frequency outliers. Second, in our neutral model, single-nucleotide polymorphisms (SNPs) are permuted based on mutation rates, and their frequencies are randomly sampled from the observed allele frequency spectrum. A more principled simulation approach that accounts for the site frequency spectrum and demography could improve the accuracy of the null distributions. Finally, although incorporating structural context provides a biologically relevant representation of the neighborhood of individual residues, HuSC only considers static structures and does not currently account for local folding dynamics or conformational flexibility, which can substantially influence the degree of constraint inferred for a given site.

Future development of this framework could incorporate an adaptive context window that uses a set of pre-defined local structural units, allowing the size of the 3D region to better reflect the local fold and differences in statistical power. Additionally, tools that generate conformational ensembles rather than static protein structures could further improve constraint estimation. The HuSC framework can also be readily extended to additional species, enabling systematic comparisons of intraspecies constraint across the tree of life and providing new insights into the evolution of mutational tolerance. Similarly, the fine-tuning strategy we introduce can be applied to more sophisticated protein language models, offering a flexible approach to integrate population-level information into increasingly powerful architectures. Together, these extensions have the potential to enhance biological resolution and provide improved tools for understanding protein evolution and interpreting variant effects.

## 4 Methods

### Mapping human variants to 3D protein structures

We followed the variant mapping approach developed in our previous COSMIS method^2^. We obtained a JSON file from github.com/capralab/cosmis that provides genetic variant counts and types (missense or synonymous) across 141,456 individuals from gnomAD v2.1.1 for all positions in the human proteome^14^. Briefly, variant statistics were extracted from GRCh38-lifted gnomAD v2.1.1. Using vcftools, sites with a FILTER flag other than PASS were removed, and only single-nucleotide variants were retained. For each human protein, we used its UniProt ID to retrieve the corresponding Ensembl transcript IDs via the UniProt ID Mapping service. Only transcripts with complete and valid coding sequences (CDSs), as confirmed by *in silico* translation, were retained. When multiple transcripts were valid, the transcript with the largest number of observed variable sites was selected.

3D protein structures were taken from the AlphaFold database version 2022-11-01^24^. For each codon, we took the minor allele frequency (MAF) of the observed variants and aggregated these values per codon position, generating dictionaries of synonymous and missense variant frequencies. 3D spatial neighborhoods for each residue were defined by parsing protein structures using Biopython’s PDBParser and computing distances between the Cα atoms for all residues. Each residue was annotated with the number of residues within a given distance (e.g., 8 Å).

### Estimating codon-specific synonymous and missense mutability

We estimated sequence context-dependent codon mutability using precomputed trinucleotide mutation rates obtained from github.com/capralab/cosmis. For each codon in the coding sequence, all possible single-nucleotide substitutions were considered. To account for local sequence effects, the substitution probability of each nucleotide was determined by the mutation rates specific to its trinucleotide context, defined as the site plus its immediately adjacent upstream and downstream nucleotides. Each potential substitution was classified as synonymous or missense, excluding substitutions that generated stop codons. For codons at the termini of the coding sequence, where only two nucleotides could be mutated within the defined context, probabilities were restricted accordingly. Mutability contributions from all three possible substitutions at each nucleotide were summed across the codon, yielding a per-codon pair of synonymous and missense mutation probabilities. This procedure was applied to all codons in the transcript, producing a nucleotide sequence context-dependent estimate of expected synonymous and missense mutability for each amino acid in the protein.

### Estimating per-transcript expected synonymous and missense variant counts

We estimated the expected number of synonymous and missense variants per transcript using a regression-based framework that models the relationship between mutability and observed variant counts under minimal selective pressure. Briefly, we computed transcript-level synonymous and missense mutability by summing codon-specific context-dependent mutabilities. We then fitted a linear regression of synonymous variant counts on synonymous mutability across transcripts, leveraging the fact that synonymous variation is largely neutral. The resulting regression model provides expected variant counts under minimal constraint given the mutability of the sequence, which we applied to per-transcript missense mutability to obtain expected missense variant counts.

### Simulating a null distribution of site-level mutation probabilities

We implemented a permutation-based simulation procedure to derive the null distribution of expected total variant frequencies for each protein site with a transcript given its expected number of missense variants. For each transcript, we normalized the missense mutability for each codon. In each permutation, we simulated *m* missense variants, where *m* is the expected count derived from the regression model described in the previous section, by sampling sites with replacement, with the probability of selecting site *i* proportional to its normalized missense mutability. At each site, a minor allele frequency (MAF) was randomly drawn from the empirical distribution of observed MAFs within the transcript and added to the site’s mutational burden. This procedure was repeated 10,000 times to generate an empirical null distribution of site-level mutational loads that follows the expected number of variants, site-wise mutation probabilities, and the variant frequency distribution.

### Calculation of Human Spatial Constraint (HuSC) scores

To compute HuSC scores for each residue within a protein, we considered its 3D spatial region. We quantified constraint by comparing the total observed frequency of missense variants in the region to a permutation-based null distribution. The observed total was calculated as the total frequency of missense variants across all residues within its 3D spatial region. To generate the null, we performed 10,000 permutations of variant placements as described above. To quantify the magnitude of deviation from the expectation, we also calculated a standardized Z-score:

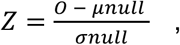

where (O) is the observed missense count, *μnull* is the mean missense count from the null distribution, and *σnull* is its standard deviation. Negative Z-scores indicate fewer observed variants than expected (constraint), whereas positive scores indicate tolerance. To improve interpretability and mitigate extreme values, we applied a log transformation, yielding the human spatial constraint Z-score (HuSC).

### Protein functional properties and other constraint scores

To evaluate the HuSC scores, we obtained a curated gene set representing different levels of essentiality from the MacArthur Lab’s *gene_lists* repository. These included haploinsufficient, essential, nonessential, autosomal dominant, autosomal recessive, and olfactory receptor genes, many of which were also used to benchmark the pLI metric. Genes were identified by HGNC symbols and mapped to UniProt accession numbers using UniProt’s API. We calculated HuSC scores for all positions for these genes. These included 141 olfactory genes, 624 non-essential genes, 904 recessive genes, 469 dominant genes, 590 essential genes, and 217 haploinsufficient genes.

### Protein Structure Visualization

Protein structures were visualized using UCSF ChimeraX (version 1.6.1)^46^. To highlight residue-specific properties, structures were colored using the B-factor replacement tool available at [Structure Coloring Studio](https://gyasu.github.io/structure-coloring-studio.html).

### Evaluations and comparisons of methods

We evaluated interspecies and intraspecies conservation/constraint metrics on the same dataset curated by our lab previously. Briefly, this dataset was retrieved from ClinVar^47^ in August 2021. The dataset only includes missense variants labeled as “Pathogenic”, “Likely pathogenic”, or “Pathogenic/Likely pathogenic” for true positive (pathogenic) variants and “Benign”, “Likely benign”, or “Benign/Likely benign” for true negative (benign) variants. Only variants with at least one-star review status and no conflicting interpretations were included. ROC and PR curves were generated and mean AUCs were reported to quantify predictive performance. For each metric (ROC AUC and PR AUC), we performed 1,000 bootstrap resamples to estimate the distribution of performance for each score. Pairwise empirical *p*-values were computed by comparing bootstrap distributions, and false discovery rate (FDR) correction was applied using the Benjamini–Hochberg method to account for multiple testing.

We compared the Spearman correlation between HuSC and several conservation metrics to assess its similarity in capturing evolutionary constraint. This dataset was curated by our lab previously. Human-variation–based metrics included RVIS^16^, missense Z^15,17^ score, pLI^15^, MTR^1^, and MTR3D^21^, while inter-species conservation metrics included GERP++^48^, phyloP^49^, phastCons^50^, and ConSurf^51^. ConSurf scores were computed using Rate4Site with default parameters and the 100-way vertebrate alignment from the UCSC Genome Browser.

### Enrichment analysis of human-specific constrained residues

To identify residues under putative human-specific constraint, we compared human spatial constraint scores (HuSC) with interspecies evolutionary conservation scores from ConSurf. We generated a scatter plot of HuSC versus ConSurf scores across residues and applied threshold criteria to define residues that are constrained in humans but not strongly conserved across species. Specifically, residues satisfying HuSC < 0 (indicative of constraint in human populations) and ConSurf > 3 (indicative of lower interspecies conservation) were classified as putatively human-specific constrained residues. While these thresholds are ad hoc, we feel this is appropriate, given that our goal was to identify a set sites and proteins most strongly enriched for constraint differences.

For each protein, we calculated the proportion of residues falling within this thresholded region. Genes were then ranked by this proportion, and the top 100 genes were selected for downstream functional enrichment analysis. Functional enrichment was performed using the Gene Ontology (GO) Biological Process annotation 2025 dataset via the Enrichr web tool^52,53^. All genes included in the HuSC–ConSurf comparison constituted the background gene set to control for dataset-specific biases.

### Missense variant scoring and protein language model evaluation

We used the ESM2 family of masked protein language models^30^. Specifically, we evaluated models with 8M, 35M, 150M, and 650M parameters. These models are trained to predict masked amino acids from sequence context and can be used to score all possible missense mutations across a protein sequence in a single forward pass.

Given a wild-type (WT) protein sequence as input, ESM2 outputs a probability distribution over the 20 standard amino acids at each residue position. Let *p(xi,a)* denote the predicted probability of amino acid *a* at position *i*. The log-likelihood ratio (LLR) score for a missense mutation from the wild-type amino acid ***WT*** to an alternative amino acid ***ALT*** at position *i* was computed as:

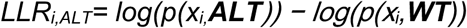

To evaluate model performance, we compared LLR-based predictions against experimentally measured variant effects from deep mutational scanning (DMS) datasets curated in the ProteinGym benchmark^32^. For each dataset, we assessed predictive performance using Spearman rank correlation between predicted LLR scores and experimental fitness measurements. In addition, we evaluated the ability of models to prioritize the most and least fit variants by computing normalized discounted cumulative gain (NDCG) for the top and bottom 10% of variants ranked by experimental fitness.

### Protein language model fine-tuning with HuSC scores

All human proteins with available HuSC scores were randomly divided into training (75%) and validation (25%) sets. We fine-tuned ESM2 models using Low-Rank Adaptation (LoRA), applied only to the self-attention modules. Specifically, LoRA adapters (rank = 8, α = 64, dropout = 0.05) were inserted into the query, key, and value projection layers of the attention blocks, following the PEFT framework (task type: *feature extraction*). LoRA introduces low-rank updates to selected layers instead of modifying all weights, substantially reducing the number of trainable parameters while preserving model capacity. This makes fine-tuning computationally efficient and memory-friendly, especially for large models.

Protein sequences were filtered to include only those with at least 25 constrained residues (HuSC < 0) and a median HuSC score < 0, ensuring an adequate signal of constraint for training. The filtered dataset was split into training (75%) and validation (25%) sets. Fine-tuning was performed with a batch size of 1 using the AdamW optimizer (learning rate = 1×10⁻⁵, weight decay = 1×10⁻²) on an NVIDIA A100 GPU.

For each batch, we computed position-specific log-likelihood ratio (LLR) scores from model logits by normalizing the wild-type amino acid probability at each position and converting these to entropy values across the 20 amino acids. The normalized entropy, representing model uncertainty, was compared to HuSC scores using a listwise ranking loss applied only to residues with negative HuSC values. To avoid overfitting, we implemented early stopping with validation checks based on mean validation loss. The model with the lowest validation loss was saved as the final fine-tuned checkpoint.

The base and fine-tuned models were evaluated using deep mutational scanning (DMS) data from the ProteinGym benchmark. We restricted the dataset to 201 proteins with sequence lengths ≤1022 amino acids to ensure compatibility with model input constraints. For each protein, log-likelihood ratio (LLR) scores were computed by passing the wild-type sequence through the model. In cases where multiple variants corresponded to the same residue, their scores were averaged to obtain a single representative value per position.

## Data availability

All datasets generated and code used in this study will be made available soon at the HuSC repository (GitHub: [github.com/gyasu/HuSC]).

