## Supplementary Figures for "Fine-tuning protein language models on human spatial constraint improves variant effect prediction by reducing wild-type sequence bias"

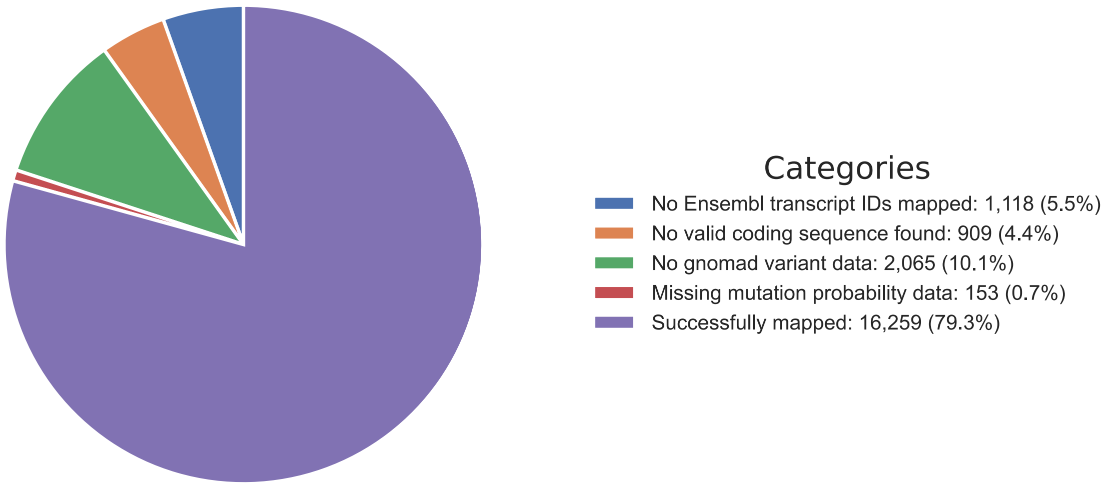


**Figure S1.** Pie chart summarizing coverage of the human proteome by the HuSC framework. HuSC scores were obtained for 16,259 proteins (79.3% of the human proteome). Proteins not covered include those lacking a mapped Ensembl ID (1,118; 5.5%), a valid coding sequence (909; 4.4%), gnomAD variant data (2,065; 10.1%), or mutational probability data (153; 0.7%). Although a protein may fail coverage for multiple reasons, only the first filtering criterion encountered is reported here.

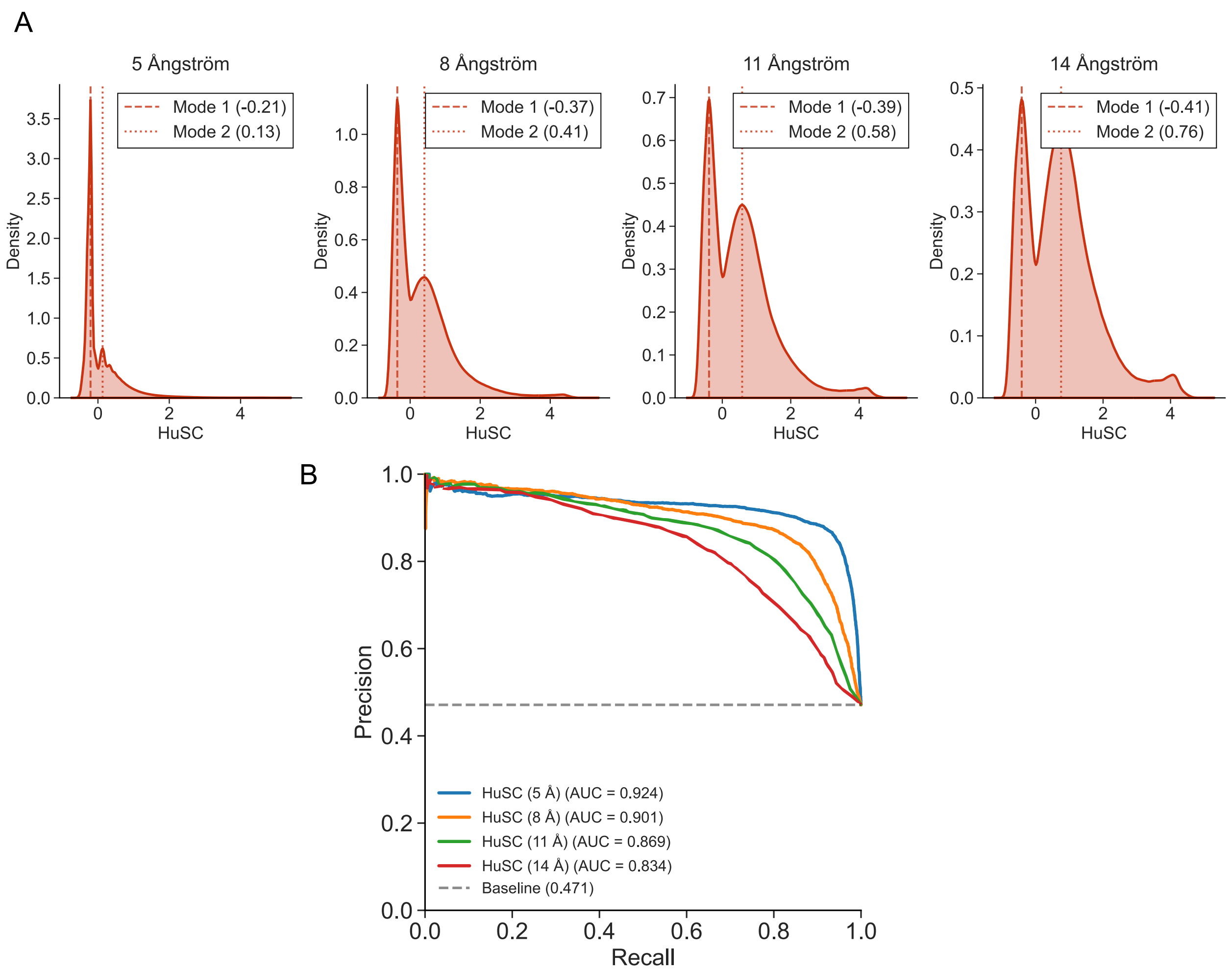


**Figure S2.** Comparison of context window sizes used for HuSC quantification. **(A)** Distribution of HuSC scores computed across 8.8 million residues from 16,259 human proteins using structural context windows of 5, 8, 11, and 14 Å. Negative HuSC scores indicate stronger evolutionary constraint, whereas positive scores indicate greater variation than expected under neutral evolution. **(B)** Precision–recall (PR) curves for pathogenicity classification using HuSC scores derived from each window size, evaluated on 6,416 pathogenic and 7,204 benign missense variants.

### Impact of the amount of spatial context on constraint quantification

To explore the influence of different amounts of 3D spatial context on constraint quantification, we applied the HuSC framework using regions ranging from 5 to 14 Å. We hypothesized that expanding the structural region would improve power to quantify constraint by capturing more sites and biologically relevant residue interactions. However, we also anticipated a trade-off between improved constraint estimation and reduced specificity when incorporating more distant residues, which could obscure constraints at smaller spatial scales.

The distributions of HuSC scores across the proteome shift systematically with increasing context region size (Fig. S2A). With increasing context region size, the HuSC distribution broadens, suggesting improved sensitivity to variation in constraint.

We investigated how well HuSC, quantified using different context region sizes, classifies pathogenic variants from ClinVar. Figure S2B shows the Precision-Recall (PR) curve for HuSC scores quantified across various context region sizes. The 5Å context regions show the highest overall PR AUC (0.92), maintaining high precision (>0.9) at high recall values. In contrast, considering more spatial context yields the highest precision in high-confidence predictions (the low-recall region of the curve), but at the cost of substantially lower precision at recall levels above 0.5.

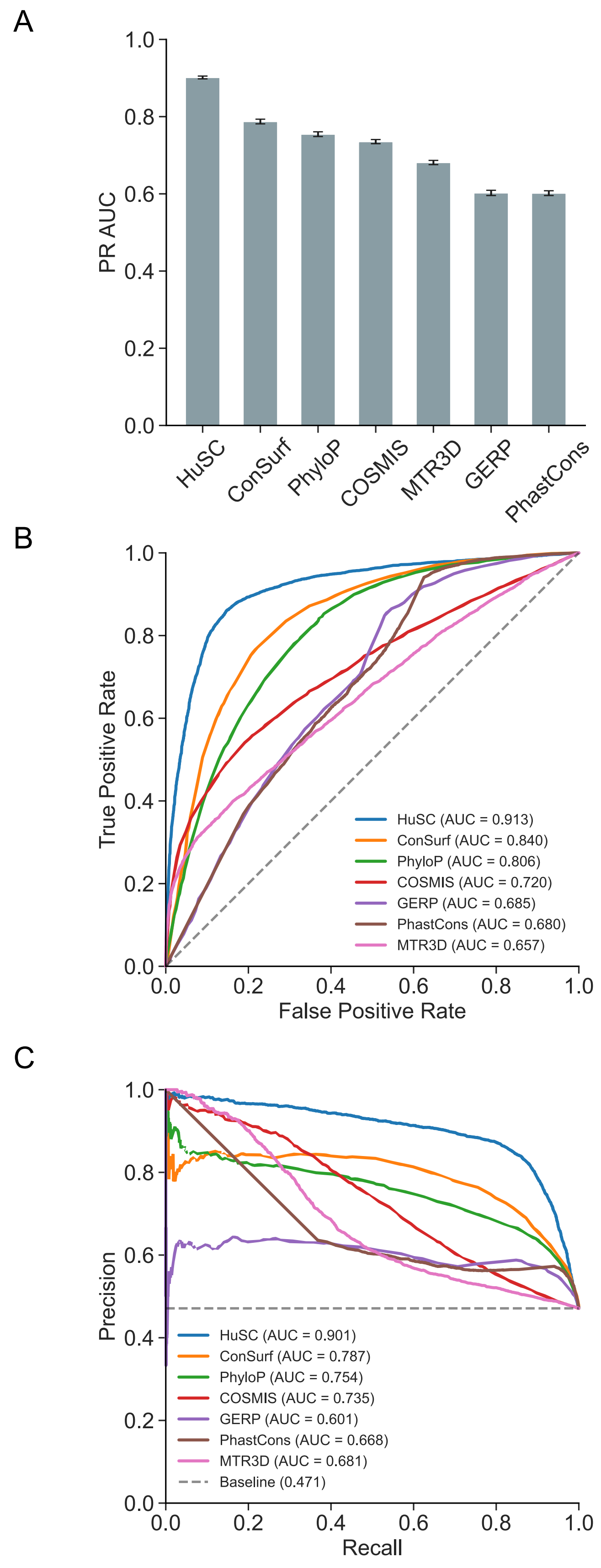
A
1.0
0.8
0.6
0.4
0.2
0.0
PR AUC
HuSC
ConSurf
PhyloP
COSMIS
MTR3D
GERP
PhastCons
B
True Positive Rate
False Positive Rate
HuSC (AUC = 0.913)
ConSurf (AUC = 0.840)
PhyloP (AUC = 0.806)
COSMIS (AUC = 0.720)
GERP (AUC = 0.685)
PhastCons (AUC = 0.680)
MTR3D (AUC = 0.657)
C
Precision
Recall
HuSC (AUC = 0.901)
ConSurf (AUC = 0.787)
PhyloP (AUC = 0.754)
COSMIS (AUC = 0.735)
GERP (AUC = 0.601)
PhastCons (AUC = 0.668)
MTR3D (AUC = 0.681)
Baseline (0.471)

**Figure S3.** Comparison of interspecies and intraspecies conservation metrics for pathogenicity classification using ClinVar variants. (A) Precision–recall (PR) area under the curve (AUC) values for classifying 6,416 pathogenic and 7,204 benign missense variants using HuSC and other conservation-based metrics. Error bars indicate the standard deviation of AUCs estimated from 1,000 bootstrap resamples of the variant dataset. Pairwise comparisons between methods were performed using the Mann–Whitney U test with false discovery rate (FDR) correction for multiple testing; HuSC achieved significantly higher performance than all other methods (FDR-corrected q < 0.05). (B) Receiver operating characteristic (ROC) curves for the same set of variants and methods. (C) Precision–recall (PR) curves for the same set of variants and methods.

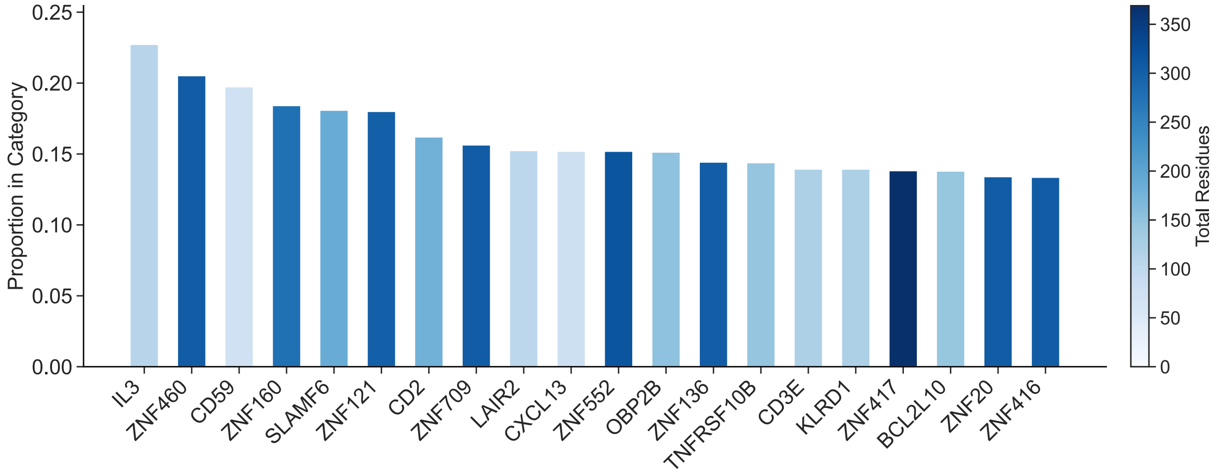


**Figure S4.** Bar graph showing genes with the highest proportion of human-specific constrained residues, as identified by HuSC. The top 20 genes are shown. Bar colors indicate the total number of residues for each gene included in the analysis.

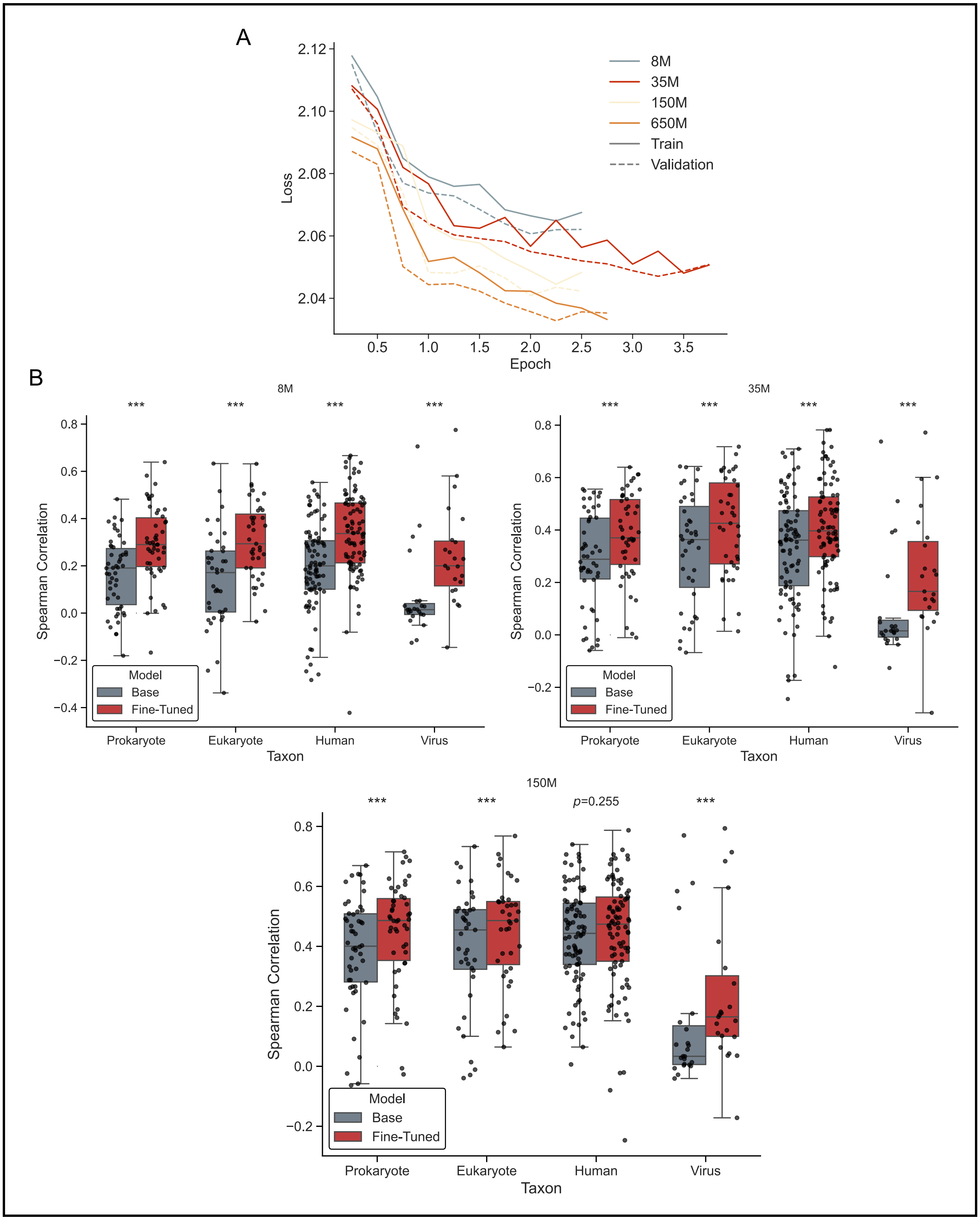
A
2.12
2.10
2.08
2.06
2.04
Loss
0.5
1.0
1.5
2.0
2.5
3.0
3.5
Epoch
8M
35M
150M
650M
Train
Validation
B
8M
35M
150M
***
p=0.255
Spearman Correlation
0.8
0.6
0.4
0.2
0.0
−0.2
−0.4
Model
Base
Fine-Tuned
Prokaryote
Eukaryote
Human
Virus
Taxon

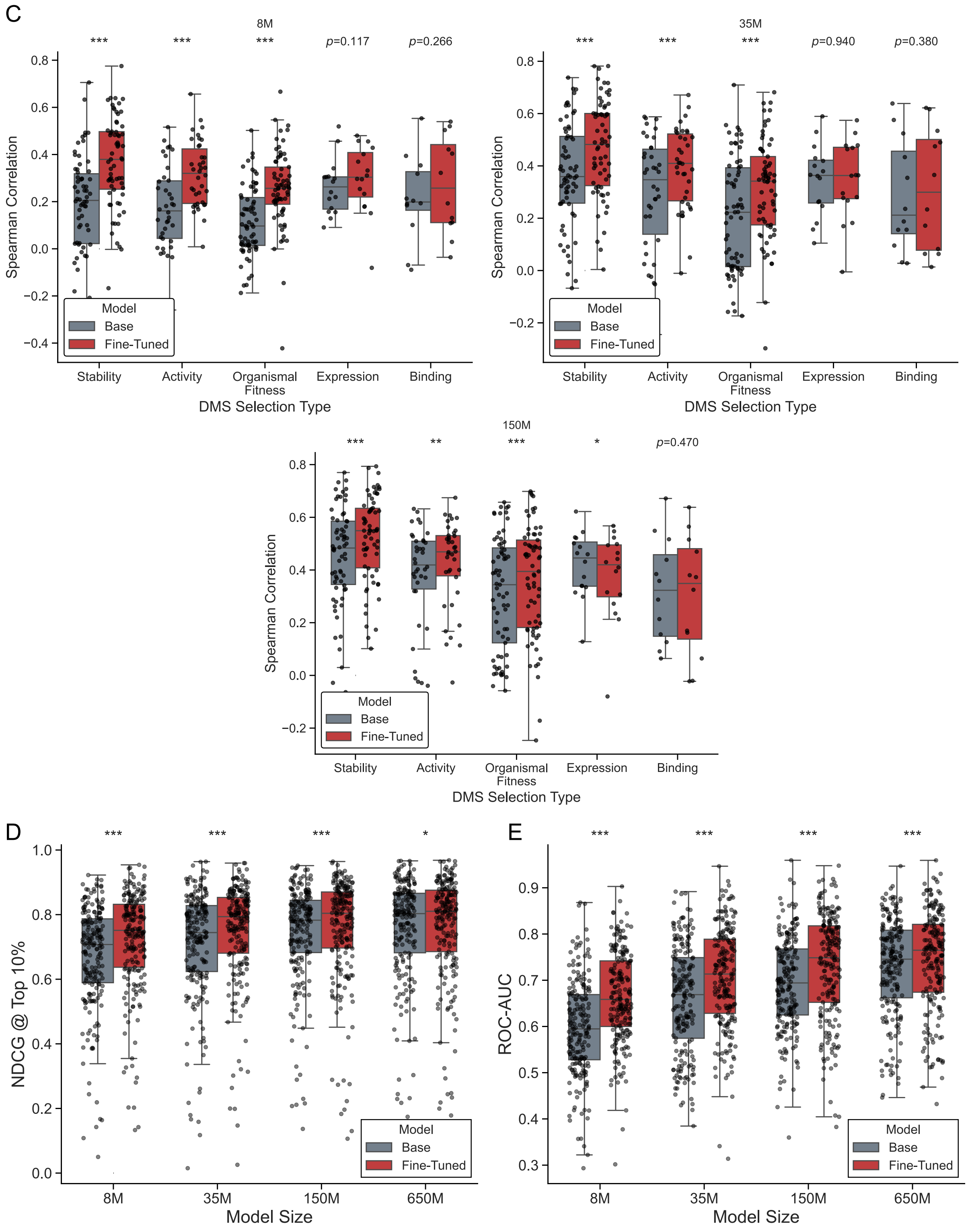


**Figure S5.** Performance evaluation of base and fine-tuned protein language models across multiple metrics. (A) Training and validation loss curves for ESM2 models with 8M, 35M, 150M, and 650M parameters. (B) Comparison of Spearman correlations between model predictions and experimentally measured deep mutational scanning (DMS) scores for 201 proteins across ESM2 models of varying sizes (8M, 35M, and 150M), before and after fine-tuning. Results are stratified by taxonomic group: 8M (left), 35M (right), and 150M (bottom). (C) Spearman correlations for the same models and protein set, stratified by DMS selection type, shown for the 8M (left), 35M (right), and 150M (bottom) models. (D) Comparison of normalized discounted cumulative gain (NDCG) at the top 10% of variants, assessing agreement between model predictions and DMS fitness scores across the same set of proteins and model sizes. (E) Comparison of receiver operating characteristic (ROC) area under the curve (AUC) values computed using binned DMS scores for the same proteins and model sizes.

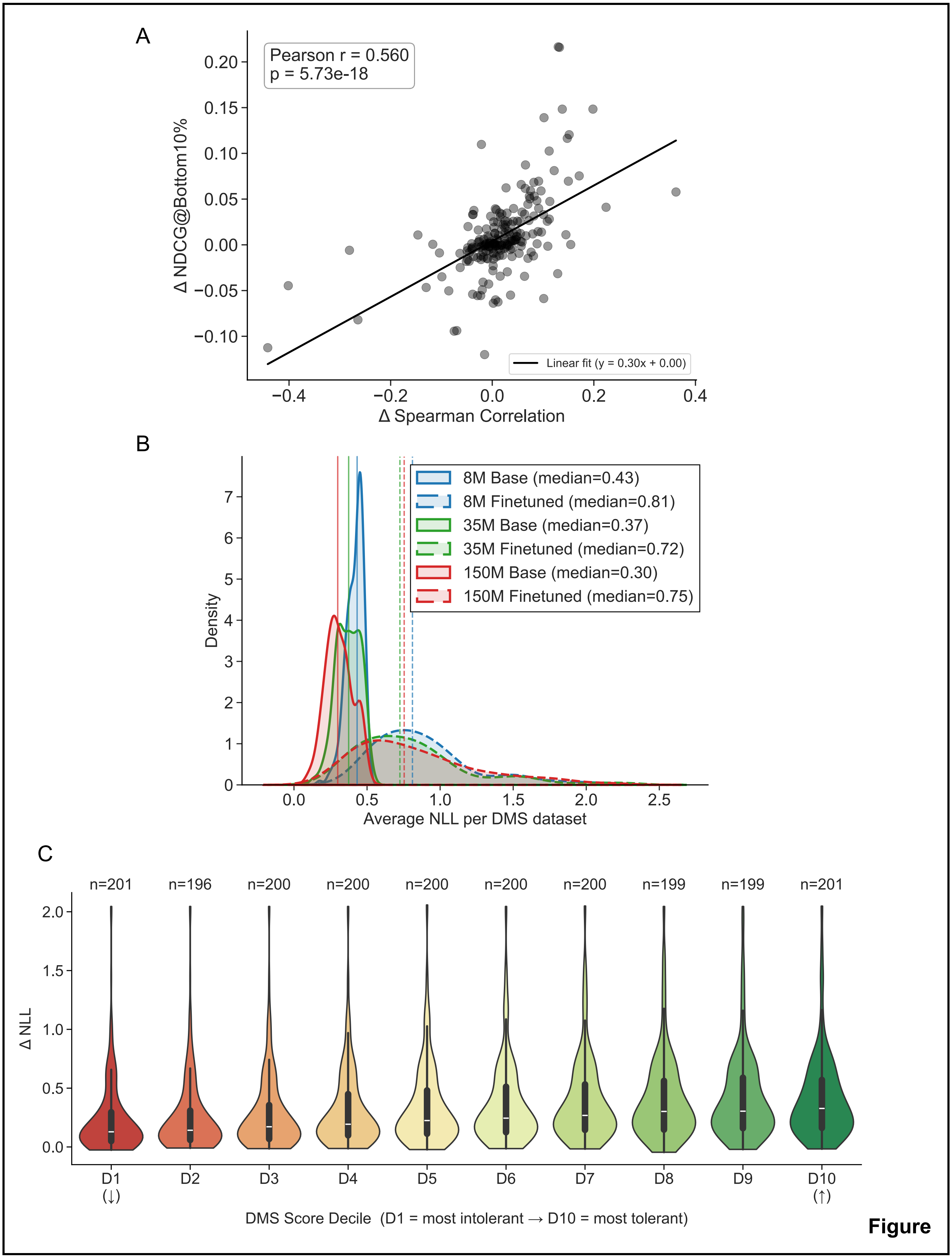
A
Pearson r = 0.560
p = 5.73e-18
Linear fit (y = 0.30x + 0.00)
Δ NDCG@Bottom10%
Δ Spearman Correlation
B
8M Base (median=0.43)
8M Finetuned (median=0.81)
35M Base (median=0.37)
35M Finetuned (median=0.72)
150M Base (median=0.30)
150M Finetuned (median=0.75)
Density
Average NLL per DMS dataset
C
n=201
n=196
n=200
n=200
n=200
n=200
n=200
n=199
n=199
n=201
Δ NLL
D1 (↓)
D2
D3
D4
D5
D6
D7
D8
D9
D10 (↑)
DMS Score Decile (D1 = most intolerant → D10 = most tolerant)

**Figure**

**S6.** Changes in model performance metrics upon fine-tuning ESM2. **(A)** Scatter plots showing the relationship between changes in model performance across all substitutions within a protein and changes in performance on the 10% least fit substitutions across deep mutational scanning (DMS) datasets for 201 proteins from the ProteinGym benchmark. Performance across all substitutions is quantified by the Spearman correlation between model predictions and experimentally measured fitness values, while performance on the least fit substitutions is measured using normalized discounted cumulative gain (NDCG). Improvements in NDCG for the bottom 10% of fitness values are strongly correlated with gains in Spearman correlation across all substitutions (Pearson $r = 0.560$, $p = 5.73 \times 10^{--18}$; linear fit: $y = 0.30x$). **(B)** Kernel density estimates of the average negative log-likelihood (NLL) of wild-type protein sequences across DMS datasets for base and fine-tuned ESM2 models with 8M, 35M, and 150M parameters. **(C)** Violin plots showing changes in NLL (ΔNLL) upon fine-tuning for residues stratified by mutational tolerance deciles, as defined by DMS fitness scores.
